# Identifying the fitness consequences of sex in complex natural environments

**DOI:** 10.1101/719252

**Authors:** Catherine A. Rushworth, Yaniv Brandvain, Tom Mitchell-Olds

**Affiliations:** Department of Evolution and Ecology, University of California, Storer Hall, One Shields Avenue Davis, CA 95616 USA; University and Jepson Herbaria, University of California, Berkeley, Berkeley, CA 94720 USA; Department of Plant and Microbial Biology, University of Minnesota, St. Paul, MN 55108 USA; Department of Biology and Center for Genomic and Computational Biology, Duke University, Durham, NC 27708 USA

**Keywords:** evolution of sex, asexuality, apomixis, reciprocal transplant, *Boechera*, hybridization, viability selection, herbivory

## Abstract

In the natural world, sex prevails, despite its costs. While much effort has been dedicated to identifying the intrinsic costs of sex (e.g. the cost of males), few studies have identified the ecological fitness consequences of sex. Furthermore, correlated biological traits that differ between sexuals and asexuals may alter these costs, or even render the typical costs of sex irrelevant. Here we use a large-scale transplant experiment of a North American wildflower to show that sex is associated with reduced lifetime fitness, despite lower herbivory. We separate the effects of sex from hybridity, finding that over-winter survival is elevated in asexuals regardless of hybridity, but herbivores target hybrid asexuals. Survival is lowest in homozygous sexual lineages, implicating inbreeding depression as a cost of sex. Our results show that the consequences of sex are shaped by complex natural environments, correlated traits, and the identity of mates, rather than sex itself.

## Introduction

Despite over a century of research, sex remains a fundamental mystery in evolutionary biology. Although recombination removes deleterious alleles and unites beneficial ones (Hill & Robertson 1966; Keightley & Otto 2006), enabling organisms to adapt (Felsenstein 1974) and evolve with antagonists (Jaenike 1978), sex has many costs. These typically include the “twofold cost,” wherein sexual organisms produce half as many offspring as asexuals due to their investment in male offspring (Maynard Smith 1978), as well as the disruption of co-adapted beneficial loci via recombination (Lehtonen *et al.* 2012). While theory has identified many costs and benefits of sex, these are likely mediated by the genomic composition of asexuals, the patterns of mating among sexuals, and the biotic and abiotic selective environment.

Despite widespread attention to the genetic processes maintaining sex, the ecological context in which sex occurs has received less attention. This is a crucial gap in our knowledge, as the processes governing sexual/asexual dynamics occur in complex natural environments. Our understanding of the ecology of sex stems largely from decades of experimental work on the freshwater snail, which has established sterilizing trematode parasites as the primary selective agent maintaining sex via frequency-dependent selection. This system offers a key example of Red Queen dynamics, wherein sexual reproduction generates novel lineages that avoid infection. Water depth moderates parasite frequency, leading to a striking ecological pattern: sex predominates in shallow water, where secondary hosts occur, while asexuality is widespread in deeper sites (Lively 1987; Jokela & Lively 1995; Lively & Dybdahl 2000; Vergara *et al.* 2014). This research shows that the abiotic and biotic environment operate in concert to shape sexual/asexual dynamics in the natural world. Critically, both ecology and its interactions with a system’s biological features provide the context necessary to pinpoint the real-life costs and benefits of sex (Neiman *et al.* 2018).

Similarly, very few studies have accounted for the widespread observation that sex and asex are often differently associated with fundamental biological and life history traits, despite the critical role these traits play in shaping the costs of sex (Meirmans *et al.* 2012). For example, the cost of males will be minimized in plants with hermaphroditic, self-compatible flowers (Charlesworth 1980; Mogie 1996); as such, frequent inbreeding will further minimize this cost. Additional costs of asexuality may be rooted in the origin and composition of asexual lineages. For example, while polyploidy and hybridity are overrepresented in asexual plants and vertebrates as compared to sexuals (e.g., Vrijenhoek 1978; Asker & Jerling 1992; Lutes *et al.* 2010), the impact of these key traits is rarely considered. Further costs of sex may be related to mate choice (inbreeding or outcrossing) and mate availability for sexual organisms, which have profound phenotypic and genetic consequences (Meirmans *et al.* 2012). Hybridization, or outcrossing between widely divergent taxa, alters numerous fitness-related traits (Yakimowski & Rieseberg 2014), which has repercussions for hybrid asexuals. The genetic impacts of mate choice will similarly influence the evolution of sex. For example, sex via inbreeding can expose a recessive genetic load (Charlesworth & Willis 2009), while asexual reproduction may maintain heterozygosity. Such life history traits likely contribute substantially to sexual/asexual dynamics, but are rarely discussed in studies of the costs and benefits of sex.

The mustard genus *Boechera* offers a rare opportunity to study the ecological costs of sex while accounting for multiple sex-associated biological traits. Apomixis (asexual reproduction via seed) is widespread in the genus (Böcher 1951; Schranz *et al.* 2005; Aliyu *et al.* 2010). These asexuals may be polyploid or diploid, enabling separate assessment of the effects of polyploidy and asexuality (Kantama *et al.* 2007; Beck *et al.* 2012; Mau *et al.* 2015). Furthermore, lineages that reproduce asexually are derived from outcrossing events, which may be either interspecific or intraspecific (Li *et al.* 2017; Rushworth *et al.* 2018), allowing separable examination of hybridity and asexuality. Finally, similar to other asexual systems (e.g., Lutes *et al.* 2010; Fradin *et al.* 2017), asexuals are highly heterozygous, whereas sexual *Boechera* are highly self-fertilizing and thus highly homozygous (Roy 1995; Song *et al.* 2006; Li *et al.* 2017). This particular trait is of broad relevance to all sexual/asexual systems, as all sexual organisms undergo shifts in population size and variable biparental inbreeding. In other words, the consequences of sex will always depend in large part on the identity of available sexual partners. For this reason, the well-characterized mating system of *Boechera* offers an ideal system for studying the evolution of sex.

Using a reciprocal transplant experiment in intact native habitat, we identified the fitness components underlying the costs of sex in *Boechera*. We examined female lifetime fitness and insect herbivory of sexual and asexual plants, finding strong over-winter viability selection against sexual lineages, which is lessened in heterozygous sexuals. This survival difference resulted in increased lifetime fitness for asexual lineages, despite elevated insect herbivory on hybrid asexuals. Importantly, the negative impacts of herbivory appeared in later life stages, reducing the probability of survival to a second year, consistent with a cost of hybridization. Numerous lines of evidence suggest that the cost of sex in this system is due at least in part to inbreeding depression in highly homozygous sexual lineages.

## Materials and Methods

### Study system

*Boechera retrofracta* (Brassicaceae) is a short-lived perennial wildflower native to western North America. Multiple *Boechera* species are highly self-fertilizing (Hamilton & Mitchell-Olds 1994; Roy 1995; Song *et al.* 2006), and *B. retrofracta*’s high microsatellite homozygosity (mean homozygosity=0.94, Rushworth *et al.* 2018) and small flowers that self-pollinate before anthesis (C. Rushworth, pers. obs.) suggest frequent self-fertilization. Nonetheless, *B. retrofracta* forms hybrids with at least 12 species (Windham & Al-Shehbaz 2006). Asexuality is tightly associated with outcrossing and hybridization in *Boechera* (Schranz *et al.* 2005; Kantama *et al.* 2007; Beck *et al.* 2012; Rushworth *et al.* 2018). Apomixis in this system is pseudogamous, requiring endosperm (but not embryo) fertilization (Böcher 1951). Because sexuals are highly selfing and asexuals must also produce a small amount of viable pollen, several of the traditional costs of sex are minimized in this system, allowing investigation of broader ecological and biological factors maintaining sex.

### Sampling and growth methods

We used 53 wild-collected diploid genotypes (24 sexual, 29 asexual) in a multi-site reciprocal transplant experiment (see Table S1 in Supporting Information). Species (or parental species, in the case of asexuals) were determined based on microsatellites and morphology (Rushworth *et al.* 2018) using the *Boechera* Microsatellite Website (Li *et al.* 2017). Heterozygosity, estimated from microsatellites, was calculated for each maternal lineage. 21 sexual genotypes were entirely homozygous, and three were not (heterozygosity ranging from 0.08 – 0.42); heterozygosity of all but one asexual genotypes was >0.5 (Figure S1). Eighteen asexual genotypes are “hybrid asexuals,” resulting from interspecific hybridization events, while eleven are “non-hybrid asexuals,” resulting from within-species outcrossing. One sexual genotype was not *B. retrofracta*, and was not analyzed; all asexual genotypes but one (“A5”) involved *B. retrofracta* as a parental species. Other parental species represented in hybrids were *Boechera sparsiflora*, *Boechera exilis*, *Boechera pendulocarpa*, *Boechera stricta*, and *Boechera puberula*. To ensure that experimental selection pressures approximated those of each genotype’s native environment, we selected lines based on proximity or similarity of collection and garden environments. All but two genotypes were collected within 10km of an experimental garden.

To minimize maternal environmental effects, genotypes were grown in controlled conditions for one generation before the experiment. Seeds were germinated on wet filter paper in petri dishes and transplanted as seedlings onto a blend of Fafard 4P Mix and MetroMix 200 (Sun Gro Horticulture, Agawam, MA, USA) in Ray Leach Cone-tainers (Steuwe and Sons, Tangent, OR, USA) in the Duke University greenhouses, and shipped as rosettes for transplanting at six weeks of age. Transplantation into replicated, randomized blocks occurred in fall for data collection the following season, allowing synchronization of flowering time with the native vegetation.

### Transplant experiment

Subsets of the 53 genotypes were planted in two cohorts of replicated, randomized blocks in the fall of 2012 and 2013. Data was collected for two growth seasons for each planting year, totaling three years of data collection. Planting year one consisted of 2000 individuals representing 19 asexual and 22 sexual genotypes, planted in five gardens with 400 plants/site (ALD, CAP, CUM, MIL, and SIL; Table S1). In year two, 2080 individuals representing 18 asexual and 21 sexual lineages were transplanted at the same sites (520 plants/site; Figure S2). However, the SIL site was excluded from year two because its dense vegetation and high humidity are atypical *B. retrofracta* habitat. Over-winter survival was scored the following spring. Plants that survived were measured for width, height, flower number, and fruit number. Due to low reproduction, MIL in year one (N=34 reproducing plants) and CUM in year two (N=2) were excluded from analyses. Leaf damage was quantified visually by trained observers, as in Prasad et al. (2012). The CUM site was excluded in herbivory analyses, as no herbivory measurements were made there. Prior to fruit dehiscence, 1 – 4 fruits were collected from 2 – 3 replicates of each genotype within each garden. Seeds were counted from each fruit and averaged across replicates to estimate average seed set per genotype per garden. Fecundity was estimated as average seed set per fruit multiplied by individual fruit number. This average was rounded to the nearest integer to allow us to model seed set as a negative binomially distributed random variable.

### Pollen viability

To evaluate how sexual system impacted pollen viability, we grew 25 sexual and 26 asexual genotypes in a growth chamber at the Duke University Phytotron. We collected mature developing buds (stage 12, Smyth *et al.* 1990) from 1 – 3 individuals per genotype (N=112 individual plants), and stained pollen to differentiate viable from inviable grains (Peterson *et al.* 2010), averaging pollen viabilities across replicates. For each individual, 100 pollen grains were randomly counted from a microscopy slide visualized on a Leica MZ7.5 Stereozoom dissecting microscope and imaged using a Canon EOS Rebel T3i digital camera.

### Statistical analyses

Our primary aim was to evaluate the association between Reproductive System (*RS*, a factor which takes the value of sexual or asexual), and group (*group*, a factor which can take the value of sexual, hybrid asexual, or non-hybrid asexual) with fitness components over time and space. Because sexuals and asexuals differ intrinsically in heterozygosity, any observed fitness differences may stem from inbreeding or outbreeding depression. We conducted additional analyses to address this possibility.

All analyses were performed using R version 3.5.2 (R Core Team 2018). Outliers were visually identified using an adjusted boxplot analysis suitable for far-right skewed data (Hubert & Vandervieren 2008), with robustbase (Maechler *et al.* 2019). To analyze fitness data, we used generalized linear mixed models (GLMMs) in glmmTMB (Brooks *et al.* 2017). Model diagnostics were conducted using DHARMa (Hartig 2019) and marginal means were estimated using the package ggeffects (Lüdecke 2018). Lifetime fitness was estimated using a zero-inflated negative binomial distribution with the canonical link functions, where structural zeros represent plants that did not survive the winter and the conditional portion of the model represents fecundity of survivors. Separate fecundity and survival models were also run, using the negative binomial and binomial error distributions, respectively. The model used was:

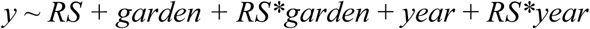

where *y* indicates a fitness component, lifetime fitness, or herbivory. Two random effects were included in all models: block nested in site and genotype nested in *RS*. The same structure was used for the zero-inflation and conditional portions of the zero-inflated negative binomial model for lifetime fitness. In all estimable models, a random effect of genotype nested in reproductive system, crossed with garden, accounted for essentially no variance, and was therefore removed from analyses. A scaled covariate for width at planting was included in survival models, while a scaled covariate of plant height was added to all herbivory models to account for a link between plant size and insect attraction.

We next tested for fitness differences between hybrid asexual genotypes, non-hybrid asexual genotypes, and sexual genotypes, by running the same models as above, replacing the *RS* term with the *group* term. To assess the effects of heterozygosity on survival, we used fixed-effects only models to analyze survival of heterozygous and homozygous sexual lineages, and complementary analyses (heterozygous sexuals vs. asexuals; homozygous sexuals vs. asexuals) were also conducted. Significance testing for fitness analyses was conducted via likelihood ratio tests (Bolker *et al.* 2009) with Holm corrections (Holm 1979).

Leaf damage was log-transformed and analyzed using a linear mixed model (LMM) in lme4 (Bates *et al.* 2015). Significance testing was conducted using *F* tests with Kenward-Roger approximation of denominator degrees of freedom (Kenward & Roger 1997) using pbkrtest (Halekoh & Højsgaard 2014).

To understand the relationship between herbivory and fitness, and how it differs among groups, two additional models were estimated with log herbivory and its interaction with group and garden as predictors. Each model examined a fitness component: fecundity and survival to a second year, both of which are conditional upon survival to year one and non-zero herbivory. The model specified was:

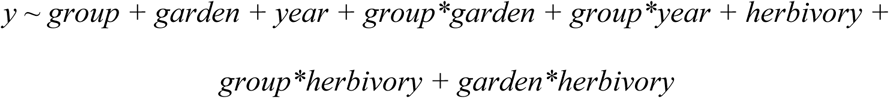

## Results

### Higher asexual lifetime fitness due to survival advantage

#### Higher asexual lifetime fitness

Lifetime fitness, the product of survival and reproduction, was higher in asexuals than sexuals (χ^2^=11.90, df=2, *P*=7.82×10^-3^). The effect of RS depended on year (χ^2^=34.74, df=2, *P*=1.72×10^-7^) and garden (χ^2^=23.45, df=8, *P*=0.016; Table S2). To disentangle the effects of hybridization and sex, we analyzed the same data with the group term. This model similarly showed an effect of group (χ^2^=13.33, df=4, *P*=0.039), and its interaction with both year (χ^2^=36.87, df=4, *P*=1.53×10^-6^) and garden (χ^2^=66.70, df=16, *P*=3.00×10^-7^; Table 1, Table S2). Estimated lifetime fitness for each group varied across year and site, but was elevated in both asexual groups, with consistently high hybrid asexual fitness (Figure S3). While our model design did not allow us to predict overall lifetime fitness for each group without accounting for interactions, raw summaries showed higher overall fitness in both groups of asexuals, although years varied slightly (Year 1 means: sexual, 211.54; hybrid asexual, 289.40; non-hybrid asexual, 299.93; Year 2 means: sexual, 76.29; hybrid asexual, 181.57; non-hybrid asexual, 145.55; Figure S4). Estimated marginal means for all models are in Appendix S1.

**Table 1.** Reproductive system (sexual vs. asexual) and hybridity predict lifetime fitness via spatially- and temporally-variable selection. Results for the main effect of group and two interactions (group and garden, and group and year) for zero-inflated negative binomial GLMMs of lifetime fitness. Left, the zero-inflation portion models structural zeros or plants that did not survive to reproduce. Center, the conditional portion models seed set in plants that survived (fecundity). Right, significance estimates for the overall model, incorporating both portions. Group indicates sexuals, “sex” vs. asexual hybrids, “hyb” vs. asexual non-hybrids, the reference category. Estimates come from conditional models, while test statistics (*χ*^2^ deviance, degrees of freedom, and *P* values) come from likelihood ratio tests for each overall effect. Significant *P*-values are in bold. Full results from each separate model are in Supplementary Information.

|  |  |  | *Zero-inflation* |  |  |  | *Conditional* |  |  |  | *Overall model* |  |  |
| --- | --- | --- | --- | --- | --- | --- | --- | --- | --- | --- | --- | --- | --- |
| term | condition | df | coef | se | $\chi^2$ | *P* | coef | se | $\chi^2$ | *P* | df | $\chi^2$ | *P* |
| Group | sex | 2 | 0.339 | 0.468 | 12.22 | **0.01** | -0.329 | 0.254 | 1.12 | 0.57 | 4 | 13.33 | **0.039** |
|  | hyb |  | -0.528 | 0.487 |  |  | -0.357 | 0.250 |  |  |  |  |  |
| Group*<br>Garden | sex*CAP | 8 | 0.129 | 0.342 | 42.82 | **6.66×10<sup>-6</sup>** | 0.048 | 0.221 | 23.70 | **0.01** | 16 | 66.70 | **3.00×10<sup>-7</sup>** |
|  | hyb*CAP |  | 0.977 | 0.349 |  |  | 0.485 | 0.208 |  |  |  |  |  |
|  | sex*CUM |  | -0.596 | 0.495 |  |  | 0.377 | 0.319 |  |  |  |  |  |
|  | hyb*CUM |  | 0.376 | 0.517 |  |  | 0.682 | 0.322 |  |  |  |  |  |
|  | sex*MIL |  | -0.278 | 0.394 |  |  | 0.052 | 0.233 |  |  |  |  |  |
|  | hyb*MIL |  | 1.063 | 0.402 |  |  | 0.435 | 0.228 |  |  |  |  |  |
|  | sex*SIL |  | -1.167 | 0.661 |  |  | 1.843 | 0.481 |  |  |  |  |  |
|  | hyb*SIL |  | -1.746 | 0.663 |  |  | 1.895 | 0.463 |  |  |  |  |  |
| Group*<br>Year | sex*2 | 2 | 1.103 | 0.378 | 23.56 | **5.36×10<sup>-5</sup>** | 0.594 | 0.194 | 13.32 | **5.16×10<sup>-3</sup>** | 4 | 36.87 | **1.53×10<sup>-6</sup>** |
|  | hyb*2 |  | -0.402 | 0.407 |  |  | 0.071 | 0.193 |  |  |  |  |  |

#### Higher asexual survival

We next asked if fitness disparities were attributable to differences in survival or fecundity. The zero-inflation portion of a zero-inflated negative binomial GLMM addresses structural zeros, or the probability of not surviving to reproduce. Over-winter survival was significantly elevated in asexual genotypes compared to sexuals. In year 1, 81% of asexuals survived, while 64.3% of sexuals survived. In year 2, 56.5% of asexuals survived, compared with only 23.8% of sexuals.

Survival differed by reproductive system (χ^2^=11.82, df=1, *P*=2.35×10^-3^), and across year (χ^2^=9.16, df=1, *P*=0.04) and garden (χ^2^=46.73, df=4, *P*=1.21×10^-8^; Table S3A). In both years asexuals had higher survival; the interactions between year and RS, and garden and RS, were also significant (year: χ^2^=22.88, df=1, *P*=8.60×10^-6^; garden: χ^2^=16.23, df=4, *P*=0.016; Table S3A). Similar results were seen in a separate model of survival (Table S4A). Plant size also increased plant survival (χ^2^=54.20, df=1, *P*=7.24×10^-13^).

Over-winter survival differed by group (χ^2^=12.22, df=2, *P*=0.01; Table 1, Table S3B), with both types of asexuals surviving better than sexuals in both cohorts (Figure 1A). The effects of group differed temporally (group × year interaction, χ^2^=23.56, df=2, *P*=5.36×10^-5^) and spatially (group × site interaction, χ^2^=42.82, df=8, *P*=6.66×10^-6^; Figure 1B, Table 1, Table S3B). A separate model of survival showed similar results (Table S4B). Hybrid asexuals were larger than other groups at planting; rosette size (χ^2^=53.79, df=1, *P*=7.24×10^-13^; Table S4B). Across gardens, hybrid asexuals had higher predicted survival than sexuals, while non-hybrid asexual survival was more variable (Figure 1B). In three of five sites, survival was roughly equivalent in both asexual groups. In two sites, survival was similarly low for sexuals and non-hybrid asexuals.

**Figure 1.**
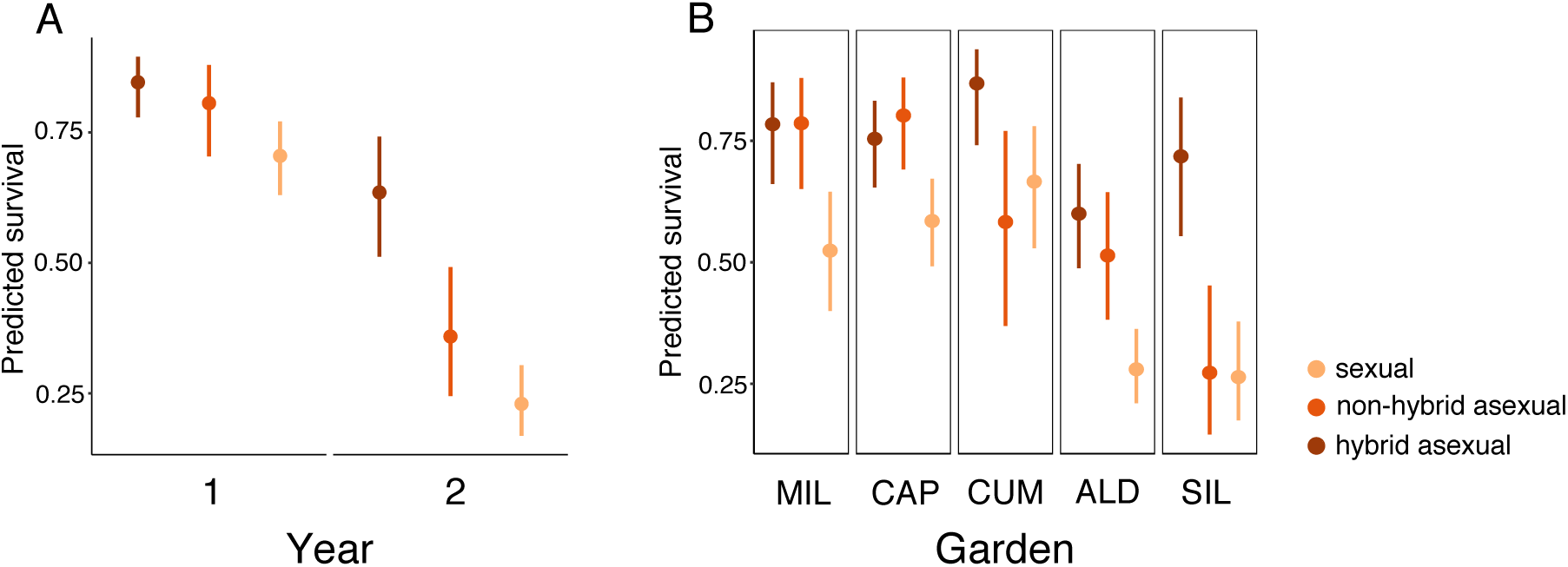
Asexuals survival is higher than sexual. Estimated marginal means for survival show substantial temporal (A, group by year interaction) and spatial variation (B, group by garden interaction). (A) Survival is reduced in sexuals in both experimental years. (B) In all sites, hybrid asexuals have higher over-winter survival than sexuals; in three of five sites, non-hybrid asexual survival is similarly high. Bars show 95% confidence intervals. SIL was only included in year one, but fitness was predicted at this site.

Heterozygosity also impacted survival. Survival of heterozygous sexuals exceeded that of homozygous sexuals (χ^2^=7.07, df=1, *P*=0.024; Figure S5, Table S5), but did not result in increased total fitness for heterozygous sexuals (Table S6). Asexual and heterozygous sexual survival did not differ (χ^2^=0.47, df=1, *P*=1). Together, these results are consistent with inbreeding depression manifesting as reduced over-winter survival in homozygous sexual lineages.

#### No differences in fecundity

We next examined the conditional portion of the zero-inflated negative binomial GLMM, which analyzed fecundity (seed set conditional on survival). In total, 995 individuals set seed (597 asexuals and 398 sexuals); removing nine outliers left a dataset of 986 individuals. Seed set did not differ intrinsically between sexuals and asexuals, but the influence of RS varied by year (χ^2^=11.86, df=1, *P*=1.72×10^-3^; Table S3A). The conditional group model produced similar results: although the main effect of group was not significant, non-hybrid asexual fecundity was lowest in both years (Figure S6A). The influence of group varied by year (χ^2^=13.32, df=2, *P*=5.13×10^-3^; Table 1, Table S3B) and by garden (χ^2^=23.70, df=8, *P*=0.01; Table 1, Table S3B, Figure S6B). There were substantial environmental effects on seed set (Table S3). Similar results were found in the separate fecundity models (Table S7). Pollen viability assays revealed no difference between sexuals and asexuals (pooled t-test, *t*=0.599 on 49 df, *P*=0.55), suggesting that any differences in seed set may be due to either maternal investment or pollen quantity. Heterozygosity had no impact on sexual fecundity (χ^2^=1.24, df=1, *P*=0.26; Table S8).

### Herbivores targeted hybrid asexuals

33% of the experiment (1379 of 4080 plants) experienced leaf herbivory. The removal of 29 high-herbivory outliers resulted in a dataset of 1350 individuals (809 asexuals, 541 sexuals). Asexual herbivory was higher than sexual, with an average of 5.45%±0.002se leaf area damaged per asexual individual (estimated marginal mean 2.14%±0.048se, 95%CI 1.72 – 2.66%). By comparison, mean sexual leaf damage across years was 3.16%±0.002se (estimated marginal mean 1.23%±0.05se, 95%CI 0.98 – 1.55%), a significant difference (*F*=22.55, df=1, 48.29, *P*=9.33e-05; Table 2A, Figure 2). Herbivory varied by garden (*F*=6.40, df=3, 62.99, *P*=2.99×10^-3^; Table 2A, Figure S7), consistent with spatial variation in herbivore communities. Herbivory was highest in hybrid asexuals (Figure 2). Group had a significant effect on herbivory (*F*=18.08, df=2, 42.39, *P*=1.26×10^-5^; Table 2B), with both hybrid and non-hybrid asexuals experiencing higher leaf herbivory than sexuals (Figure 2). Plant height was significant in both models (Table 2).

**Figure 2.**
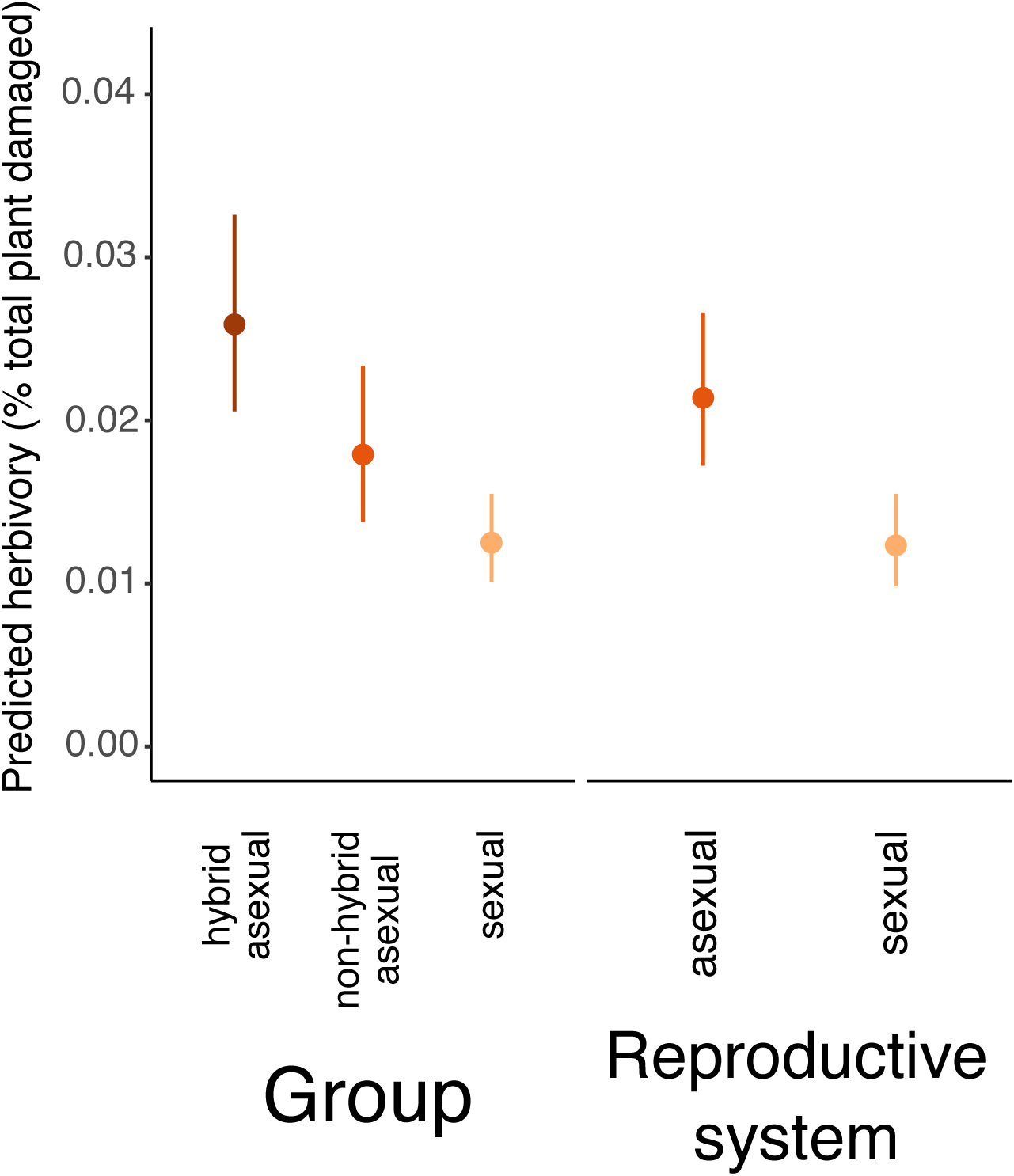
Herbivory is elevated in hybrid asexuals. Estimated marginal means for herbivory from LMMs show the main effects of group (sexual vs. non-hybrid asexual vs. hybrid asexual, left) and reproductive system (sexual vs. asexual, right). Bars show 95% confidence intervals.

**Table 2.** Herbivory varies by reproductive system, hybridity, and garden. (A) Model results for reproductive system (“RS”, sexual vs. asexual) LMM and (B) group (sexual “sex” vs. asexual hybrid “hyb” vs. asexual non-hybrid, the reference category) LMM. Significant *P*-values are in bold. Estimates apply to the conditional models, while test statistics (*F* statistics, degrees of freedom, and *P* values) come from significance tests for each overall effect.

| term | condition | estimate | se | df | *F* | *P* |
| --- | --- | --- | --- | --- | --- | --- |
|  | intercept | -1.633 | 0.071 | - | - | - |
| RS | sexual | -0.299 | 0.079 | 1, 48.29 | 22.55 | **9.33×10<sup>-5</sup>** |
| Garden | CAP | -0.002 | 0.074 | 3, 62.99 | 6.40 | **2.99×10<sup>-3</sup>** |
|  | MIL | 0.340 | 0.095 |  |  |  |
|  | SIL | -0.002 | 0.103 |  |  |  |
| RS*Garden | sexual*CAP | 0.049 | 0.080 | 3, 1280.95 | 1.74 | 0.372 |
|  | sexual*MIL | 0.059 | 0.115 |  |  |  |
|  | sexual*SIL | 0.234 | 0.105 |  |  |  |
| Year | 2 | -0.045 | 0.076 | 1, 67.59 | 0.38 | 1 |
| RS*Year | sexual*2 | -0.013 | 0.084 | 1, 1175.63 | 0.02 | 0.876 |
| Plant height | - | 0.045 | 0.016 | 1, 1318.25 | 7.52 | **0.019** |

B.
| term | condition | estimate | se | df | *F* | *P* |
| --- | --- | --- | --- | --- | --- | --- |
|  | intercept | -1.719 | 0.095 | - | - | - |
| Group | sex | -0.210 | 0.010 | 2, 42.39 | 18.08 | **1.26×10<sup>-5</sup>** |
|  | hyb | 0.141 | 0.102 |  |  |  |
| Garden | CAP | -0.035 | 0.098 | 3, 62.98 | 6.37 | **2.99×10<sup>-3</sup>** |
|  | MIL | 0.332 | 0.122 |  |  |  |
|  | SIL | 0.239 | 0.149 |  |  |  |
| Group*Garden | CAP*sex | 0.083 | 0.102 | 6, 1273.93 | 1.99 | 0.065 |
|  | CAP*hybrid | 0.060 | 0.102 |  |  |  |
|  | MIL*sex | 0.067 | 0.139 |  |  |  |
|  | MIL*hybrid | 0.016 | 0.132 |  |  |  |
|  | SIL*sex | -0.005 | 0.150 |  |  |  |
|  | SIL*hybrid | -0.324 | 0.152 |  |  |  |
| Year | 2 | -0.087 | 0.099 | 1, 66.98 | 0.30 | 1 |
| Group*Year | sex*2 | 0.023 | 0.105 | 2, 1056.46 | 0.45 | 0.77 |
|  | hybrid*2 | 0.092 | 0.105 |  |  |  |
| Plant height | - | 0.042 | 0.016 | 1, 1320.42 | 6.96 | **0.019** |

### Herbivory negatively impacted survival to subsequent years

What are the effects of herbivory on fitness, and do these effects differ between sexuals and asexuals? We next predicted two separate fitness components, fecundity and survival to a second year, using models incorporating a main effect of log-transformed herbivory, and its interaction with both group and garden.

Over 90% of experimental fitness occurred in the first year of life. In both years, more asexuals than sexuals survived to a second year. In experimental year 1, 8.2% of plants survived for a second year (N=163: 108 asexuals, 55 sexuals). Similarly, 5.9% of plants in year 2 survived for a second year (N=123: 104 asexuals, 19 sexuals). Roughly 1% of plants reproduced in their second year (Year 1: 1.4%, N=28: 24 asexuals, 4 sexuals; Year 2: 1.3%, N=26: 25 asexuals, 1 sexual). Nevertheless, survival to subsequent years could substantially increase lifetime fitness.

In temperate montane environments, herbivory and seed set co-occur during the short growing season, and may thus influence one another. We found no effect of herbivory on fecundity (χ^2^=0.28, df=1, *P*=0.598; Table S9), but herbivory decreased survival to a second year (χ^2^=21.26, df=1, *P*=8.02×10^-6^; Table S10). Group also affected subsequent survival (χ^2^=32.88, df=2, *P*=5.79×10^-7^; Table S10), with more hybrid asexuals than non-hybrids or sexuals surviving (Year 1: 82 hybrid asexuals, 17 non-hybrid asexuals, 41 sexuals; Year 2: 75 hybrid asexuals, 28 non-hybrid asexuals, 17 sexuals). In all groups, herbivory and probability of survival were negatively correlated (Figure 3). Although the interaction between group and herbivory was non-significant, the negative relationship was strongest in hybrid asexuals (Figure 3), consistent with herbivory limiting hybrid asexual fitness.

**Figure 3.**
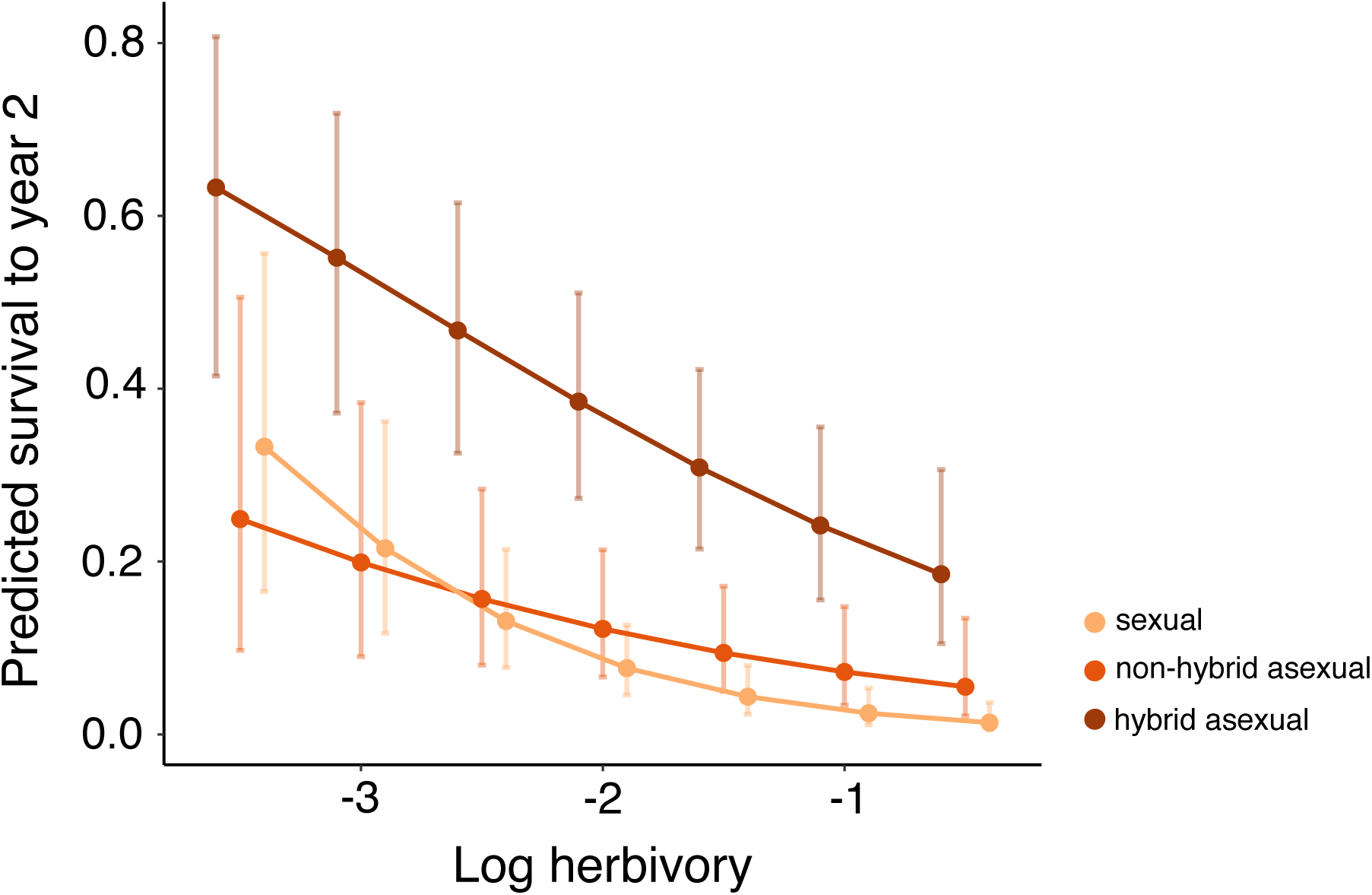
Plants with less herbivory are more likely to survive to the next year. Herbivory has a significant impact on survival to a second year, suggesting that the detrimental consequences of herbivory may not be observed in a single growing season. Although the correlation between year 2 survival and herbivory differs qualitatively by group (sexual vs. non-hybrid asexual vs. hybrid asexual), the interaction is not significant. Estimated marginal means are shown averaged across five different values of log herbivory. Points are jittered for ease of reading. Bars show 95% confidence intervals.

## Discussion

Here we show that both abiotic and biotic factors play key roles in the maintenance of sex in nature. While viability selection favors both inter- and intraspecific asexuals, implicating shared causative traits such as asexuality or heterozygosity, biotic interactions are especially influenced by hybridization. Together, these results suggest that, although survival of adverse abiotic conditions is facilitated by asexuality or associated heterozygosity, hybridization plays a singular role in the interaction between asexuality and herbivory. The precise targets of selection are currently unknown, but our results suggest that the costs of sex depends on ones’ sexual partners, and may be mediated by interactions with biological antagonists.

### The complex ecology of sex

Despite extensive theoretical work and recurrent calls for field-based studies (Agrawal 2006, 2009), few researchers have examined the ecology of sex. Identifying ecologically-relevant traits associated with sex, and the selective processes acting in natural environments, could elucidate the phenotypes and ultimately the molecular mechanisms underlying the cost of sex in wild populations.

Although numerous selective agents are likely important to the evolution of sex (Neiman *et al.* 2018), we have little understanding of how these biotic and abiotic selection pressures maintain sex in the wild. Previous work shows that sex may permit survival during abiotic environmental shifts (Stross & Hill 1965) or enable avoidance of fitness-reducing biotic interactions (Lively 1987). We found that strong viability selection early in life results in higher lifetime fitness for asexual lineages, but increased insect herbivory may provide an upper limit on asexual fitness. That different agents act at different life stages suggests that long-term studies are key to understanding the evolution of sex.

Many of the best-known asexual systems are polyploid (e.g., Fox *et al.* 1996; Thompson & Whitton 2006), hybrid (e.g., Xu *et al.* 2015) or both (e.g., Lutes *et al.* 2010; Gutekunst *et al.* 2018). Yet these biological features are rarely considered in either studies of these systems or in broader discussions of the costs of sex. We explicitly considered traits correlated with reproductive mode in assessing patterns of selection on sexual reproduction, finding that viability is especially reduced in homozygous sexual lineages, consistent with inbreeding depression acting as a cost of sex. Hybrid asexuals were more prone to herbivory, suggesting that the costs of asexuality may be related to the correlated trait of hybridity. Clearly, selective processes that influence the evolution of sex are shaped by both the origin and composition of asexual lineages and patterns of mating in sexuals.

### Mating system influences the evolution and ecology of sex

The fundamental role of sex in all organisms is to reshuffle variation across the genome through recombination and segregation. But the effects of these processes are tempered by the efficacy of such shuffling. When sexual partners resemble one another due to population structure, limited population genomic variation and/or inbreeding, the traditional costs and benefits of sex can change dramatically (De Visser & Elena 2007). For example, self-fertilization cannot unite beneficial mutations arising in different lineages. Practically speaking, the costs and benefits of sex are always a function of mate availability and identity.

Despite the widespread existence of asexual reproduction in flowering plants, little is known about the costs of sex in angiosperms. Researchers studying the evolution of sex in natural populations have largely focused on animal systems, where the existence of separate sexes provide clear predictions about the relevant costs and benefits (Lehtonen *et al.* 2012). Although hermaphroditic asexuals are still predicted to have a 1.5-fold fitness advantage, sex via self-fertilization muddles these predictions (Charlesworth 1980), as do additional biological traits common in plants (Mogie 1996). For example, both selfing and pseudogamous apomictic *Boechera* must produce sufficient pollen for self-fertilization, reducing the applicability of any cost of males. Additionally, the strong link between hybridization and asexuality in numerous vertebrate and plant taxa (Asker & Jerling 1992; Kearney 2005) suggests that recurrent independent hybridization events generate novel asexuals. In *Boechera*, both sexual and asexual lineages have limited variation, but high diversity among asexual lineages may be generated by multiple independent origins of asexuality. Such cases require consideration of additional costs and benefits of sex absent from most models.

We hypothesize that inbreeding depression manifests as the primary cost of sex in sexual *Boechera*. Our experiment showed fitness costs and benefits associated with sex that follow predictions for inbreeding depression. Sexuals had reduced overwinter survival (Figure 1), particularly in homozygous lineages (Figure S4), that varied both spatially and temporally (Figure 1, Table 1, Table S4), consistent with inbreeding depression (Armbruster & Reed 2005; Cheptou & Donohue 2010). Our experiment contained few heterozygous sexuals, and was thus underpowered to detect a strong relationship between heterozygosity and fitness. Nonetheless, our results suggest that the main cost of sex in *Boechera* may be related to deleterious load, and that outcrossing may be sufficient to increase survival in this system. This pattern, known as a heterozygosity-fitness correlation or HFC, is widespread in wild systems (Szulkin *et al.* 2010). Despite reduced survival in homozygous sexual lineages, we saw no evidence that survival differed between asexuals and heterozygous sexuals (Table S5), consistent with the hypothesis that the benefit of heterozygosity is the masking of genome-wide recessive load.

Although inbreeding likely poses a main cost of sex in this system, other factors may also be at play. First, asexual lineages likely arise frequently and those found in nature have survived selection, potentially resulting in an overrepresentation of high-fitness asexual genotypes in the field. Indeed, the majority of de novo *Boechera* hybrids have lower fitness than their selfed progenitors (Rushworth & Mitchell-Olds 2020), suggesting that fitness variation may be more common in the wild than our experiment suggests. Additionally, the recurrent origins of asexual lineages (Sharbel & Mitchell-Olds 2001) suggest that asexuals comprise a range of ages. With deleterious mutations accumulating over the lifespan of a given asexual lineage, older lineages may have reduced fitness. Lastly, reproductive isolation that varies across populations may also result in fitness that depends on interactions between parental genotypes. Each of these scenarios is plausible and provides avenues for further research.

Self-fertilization also has clear benefits. Sex and recombination enable adaptation to novel environments (Felsenstein 1974), and inbreeding may speed local adaptation. In situations where environmental changes are substantial, fixation of beneficial recessive alleles through self-fertilization results in rapid adaptation (Glémin & Ronfort 2012). Self-fertilization will also reduce the potential for reshuffling of beneficial allelic combinations, as high homozygosity limits opportunities for functional disruption (Hartfield *et al.* 2017). Our results showed that sexual lineages experienced less insect herbivory than asexuals (Figure 2). Local adaptation underlies variation in chemical defense both among populations of *Boechera stricta* (Prasad *et al.* 2012) and between species (Windsor *et al.* 2005). We thus hypothesize that interactions between divergent alleles in outcrossed asexuals disrupt chemical defenses, consistent with outbreeding depression.

### Herbivory as an ecological cost of asexuality

Classic literature posits multiple costs of asexuality, including the accumulation of deleterious mutations and the extinction of asexual clones with the highest mutation load (Müller 1964; Felsenstein 1974). While asexual *Boechera* do show elevated putatively deleterious mutations (Lovell *et al.* 2017), the heterozygous state of these mutations, the higher fitness of asexuals observed here, and the replenishment of asexuals by novel hybridization events, suggests that these intrinsic genetic costs have relatively little to do with the costs of asexuality in *Boechera*.

Rather, we found evidence for a short-term ecological cost associated with asexuality in this system – a 73% increase in asexual insect herbivory (Figure 2), consistent with predictions from theoretical literature (Stearns 1987). This pattern may be underlain by asexuality, heterozygosity, hybridization, or some combination of all three. In functionally asexual *Oenothera*, generalist herbivory is elevated across multiple species (Johnson *et al.* 2009), supporting a role for asexuality in the macroevolution of plant defense. Microevolutionary patterns suggest the same; viral infection is elevated in asexual and common sexual grass genotypes (Kelley 1994). Although elevated in both asexual groups, insect herbivory is highest in hybrid asexuals (Figure 2). Ample research also shows that hybridization strongly alters plant defense and herbivore preference (Strauss 1994; Fritz 1999; Whitham *et al.* 1999; Orians 2000), and hybrid taxa may produce greater or lesser quantities of chemical compounds, and/or novel ineffective compounds (Orians 2000). Disentangling the influence of asexuality and hybridization on defense offers a fruitful avenue for future research.

While the costs of herbivory were insufficient to overcome the other benefits of asexuality in our two-year study, these costs could manifest in other traits or at later life stages (Marquis 1992). We found that herbivory decreased survival to subsequent years (Figure 3), consistent with the negative correlation between fitness and herbivory documented in *Boechera stricta* (Prasad *et al.* 2012). Similar patterns have been identified in animals, suggesting that downstream consequences of parasitism or herbivory are broadly applicable. One recent study of Soay sheep showed that fecundity and parasite load collectively reduce overwinter survival (Leivesley *et al.* 2019). Additionally, we found that the relationship between second-year survival and herbivory is strongest in hybrid asexuals, suggesting that hybridization may limit the spread of asexuality via herbivory. However, more hybrids survive to a second year of life overall. Further work, including herbivore exposure experiments, are needed to understand the interplay between fitness and herbivory in this system.

Collectively, the work presented here shows that sexual/asexual dynamics in wild plant populations are largely shaped by environmental complexity and correlated traits, including mating system. Asexual *Boechera* have consistently higher fitness and higher herbivory damage than self-fertilizing sexual lineages, and survival is lowest in homozygous sexuals. Our results highlight the importance of both the biotic and abiotic environment in the evolution of sex, and suggest that inbreeding and outbreeding depression underlie the ecological costs and benefits of sex in this system. Collectively, this work shows that the complex natural environment and the choice of a sexual partner work in tandem to shape patterns of reproductive variation in the wild.

## Acknowledgements

The authors wish to thank Sailee Clemens, Rose Keith, Cheng-Ruei Lee, Marshall McMunn, Tim Park, Katie Putney, Evan Raskin, Kara Stiff, Chris Strock, Maggie Wagner, and Ash Zemenick for their help with field work. We are especially grateful to Rui-Min Diana Mao for her work quantifying seed set and pollen viability. We also thank Julius Mojica for his statistical expertise and Rob Colautti for his help with both statistics and in the field, and John Willis, Mohamed Noor, Mark Rausher, Britt Koskella, and Michael Windham for advice. This work was supported by funding from the National Science Foundation (Graduate Research Fellowship to CAR and DEB-1311269 to CAR and TM-O) and the National Institutes for Health (R01 GM086496 to TM-O).

