## Supplementary material for "Identifying the fitness consequences of sex in complex natural environments": Figure S1

**Figure S1. Histogram of heterozygosity for maternal genotypes.** Heterozygosity is higher in asexual than sexual lineages. Three sexual genotypes used in this experiment had heterozygosity above zero.

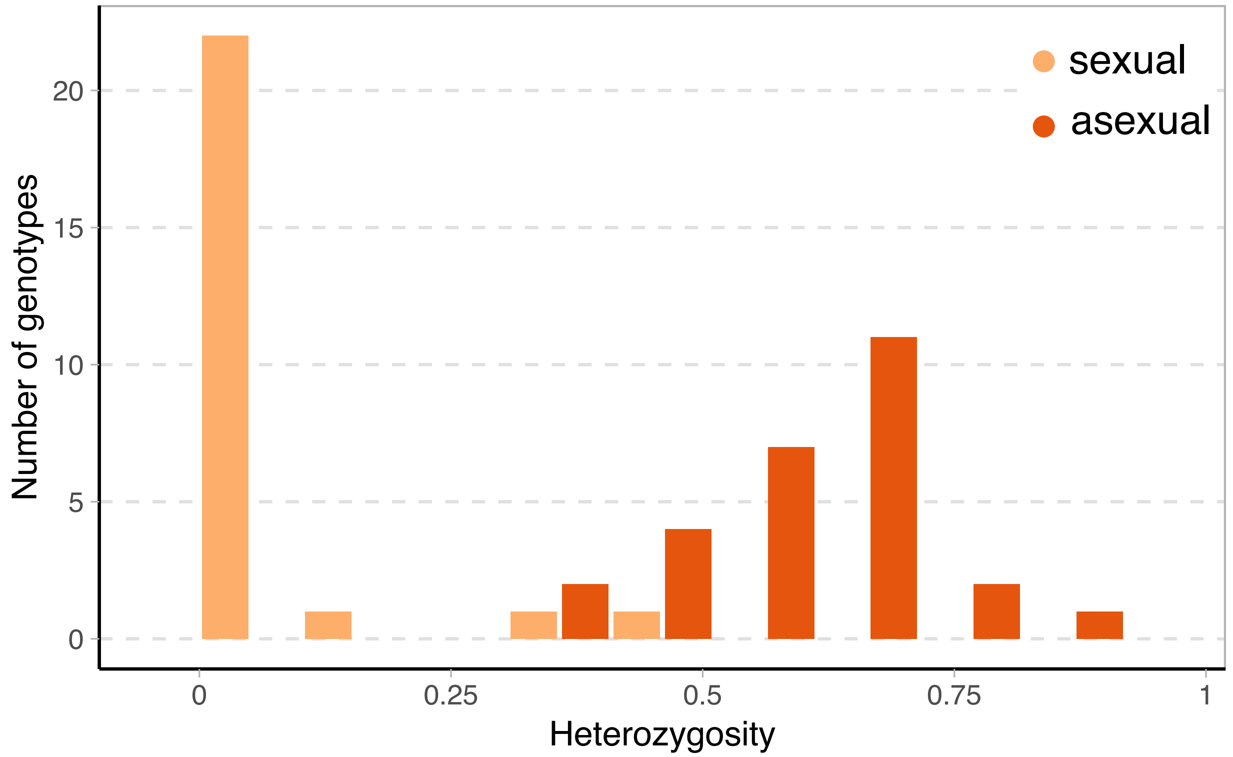
