## Supplementary material for "Identifying the fitness consequences of sex in complex natural environments": Figure S2

**Figure S2**. Map of experimental study location and garden sites in central Idaho, USA. The SIL site was only used in year 1, indicated by the light orange marker.

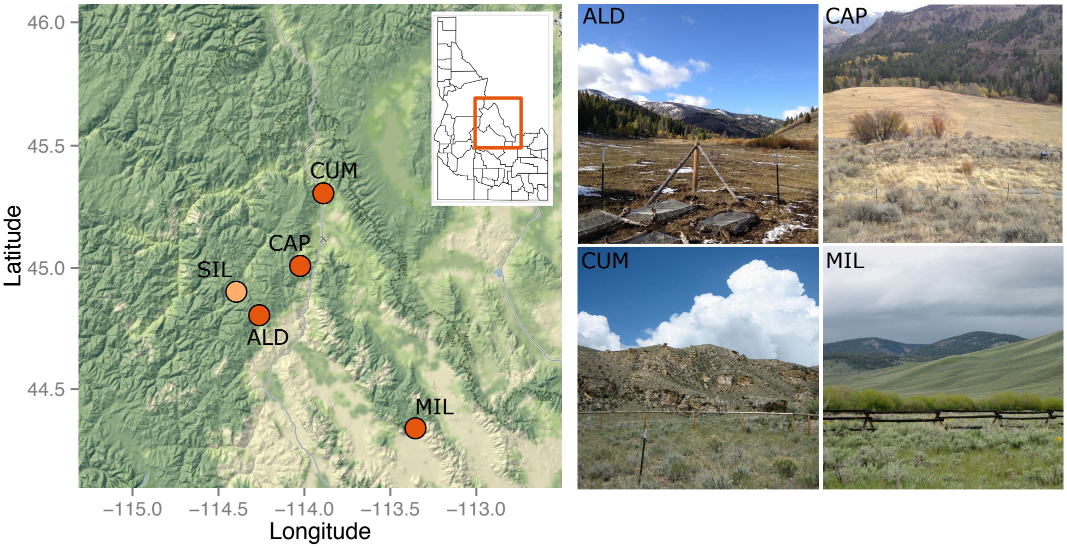
