## Supplementary material for "Identifying the fitness consequences of sex in complex natural environments": Figure S3

**Figure S3. Predicted lifetime fitness of asexuals is higher than that of sexuals.** Estimated marginal means for lifetime fitness (composed of survival and fecundity) on the log scale by group across experimental years (Year 1, top row; Year 2, bottom row) and gardens (left to right). Fitness is consistently elevated in hybrid asexuals, with non-hybrid asexual fitness often overlapping. Note that SIL was not planted in year 2, but predicted fitness from the model is shown. Bars show 95% confidence intervals.

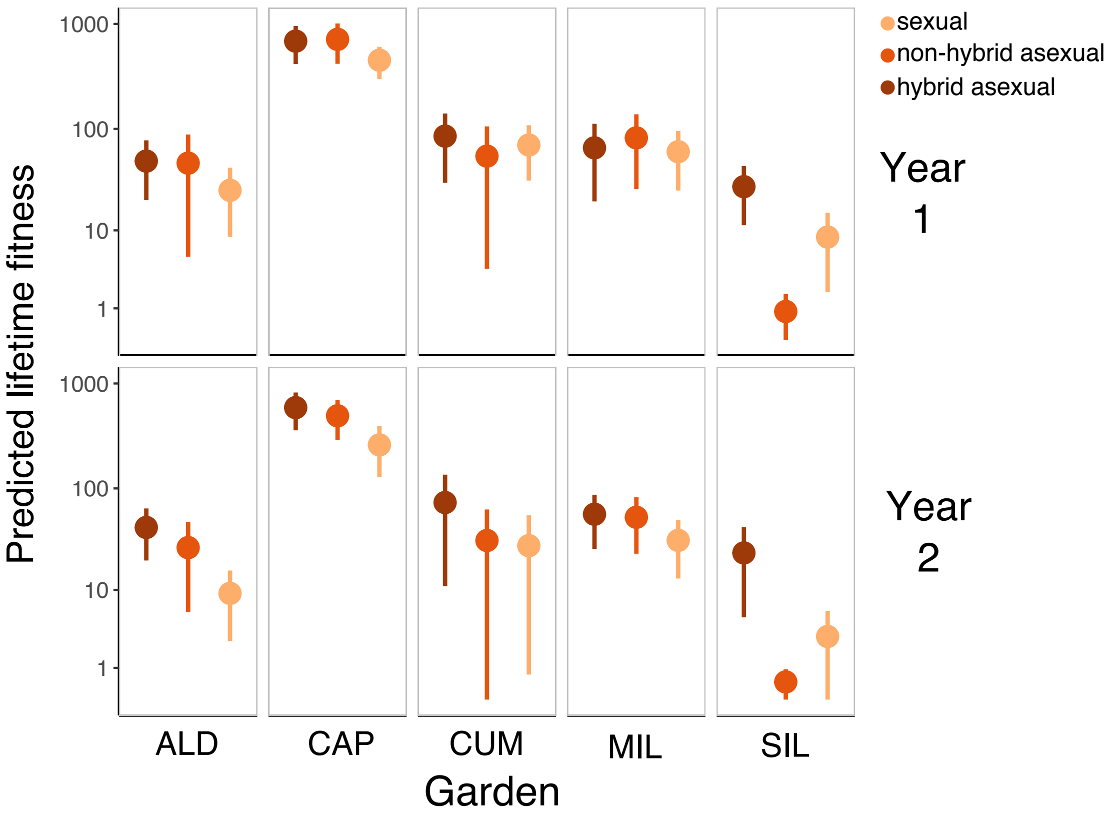
