## Supplementary material for "Identifying the fitness consequences of sex in complex natural environments": Figure S4

**Figure S4. Raw data show lifetime fitness of asexuals is higher than sexuals.** In year 1, both groups of asexuals have higher fitness. In year 2, hybrid asexuals have highest fitness, but non-hybrid asexual fitness is strongly overlapping. Solid lines indicate medians, while dotted lines indicate means.

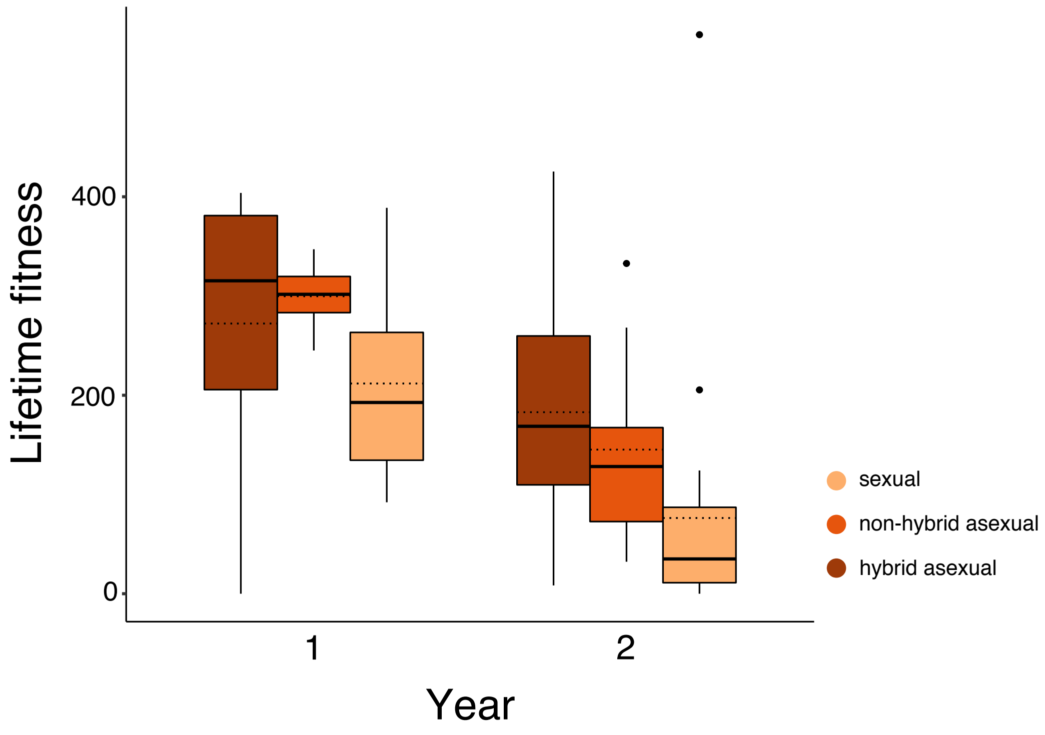
