## Supplementary material for "Identifying the fitness consequences of sex in complex natural environments": Figure S5

**Figure S5. Heterozygous sexuals have higher over-winter survival than homozygous sexuals.** This result is consistent in both experimental years. Estimated marginal means are shown; bars show 95% confidence intervals.

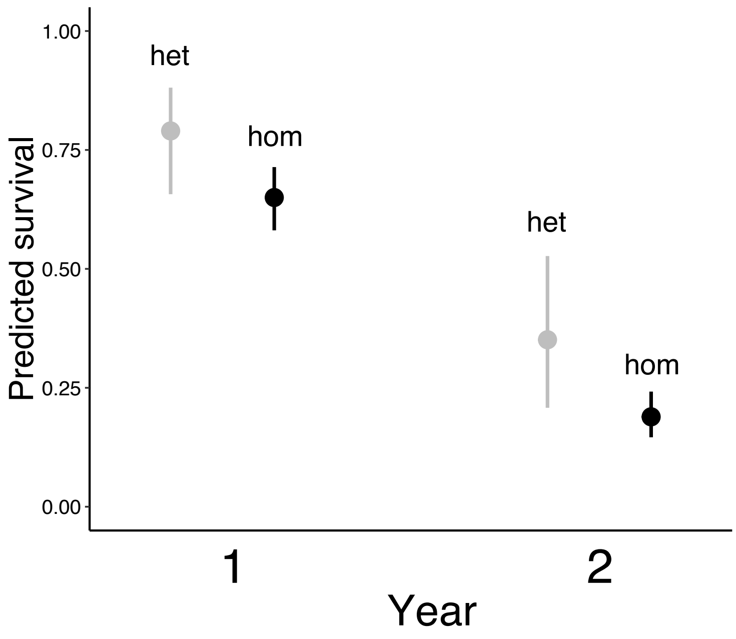
