## Supplementary material for "Identifying the fitness consequences of sex in complex natural environments": Figure S6

**Figure S6. Temporally and spatially-variable fecundity selection.** (A) Fecundity varies temporally, with higher asexual fecundity in year 1 and higher sexual fecundity in year 2. (B) Sexual fecundity (shown on log scale) is higher than, or equivalent to, asexual in some garden sites. Bars show 95% confidence intervals.

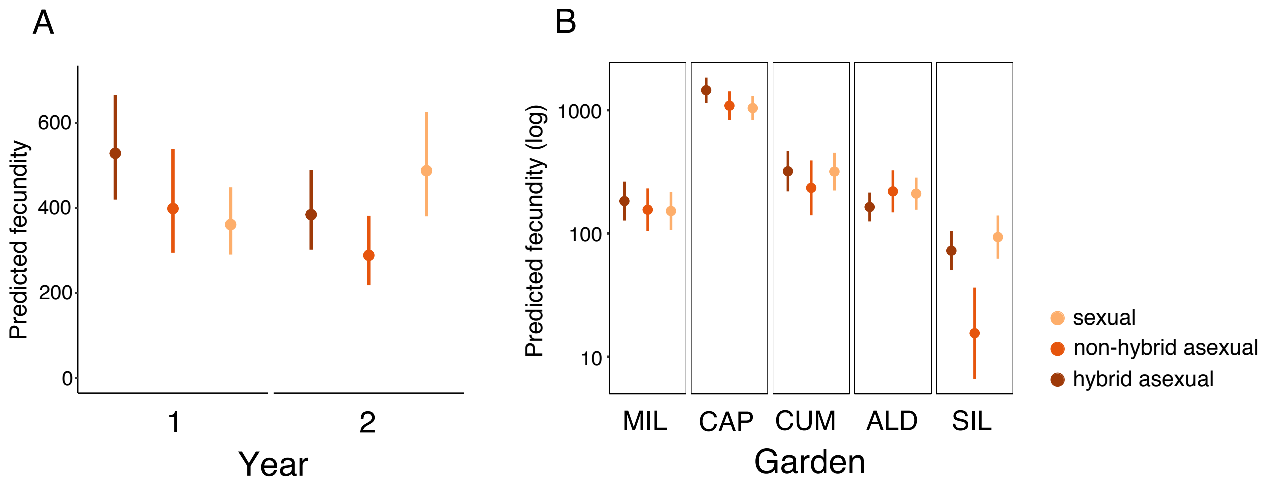
