## Supplementary material for "Identifying the fitness consequences of sex in complex natural environments": Figure S7

**Figure S7**. **Herbivory is spatially variable.** Estimated marginal means from herbivory LMM, adjusted for plant height of 0cm. Bars show 95% confidence intervals.

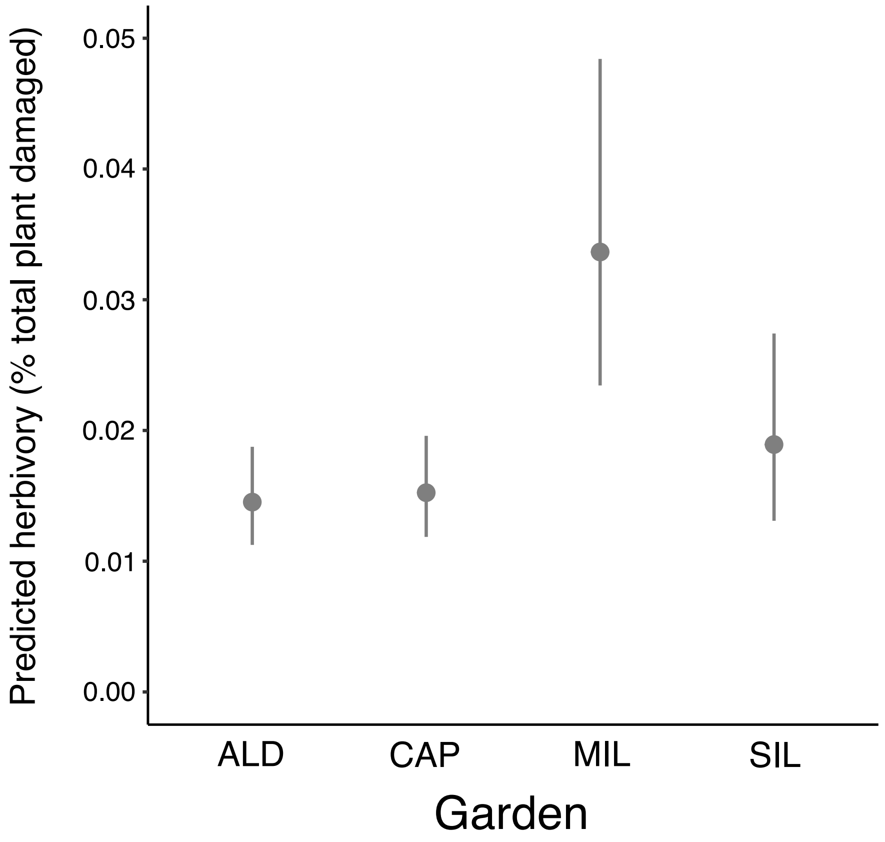
